# A crypt-operating DNA repair checkpoint for uncoupling regeneration and tumorigenesis

**DOI:** 10.64898/2026.02.27.708480

**Authors:** R. Ruez, K. Radulovic, JJ. Martinez-Torres, O. Boulard, Mound. Abdallah, T. Paz Del Socorro, G. Nigro, M. Seillier-Turini, G. Muharram, JJ. Martinez-Garcia, F. Gerbe, E. Floquet, I. Van Seuningen, A. Carrier, P. Jay, T. Sanmuganantham, ANR. Wewer, A. Försti, C. Cochet, A. Vincent, P. Rosenstiel, C. Abbadie, I. Sobhani, O. Filhol-Cochet, N De Oliveira Alves, M Chamaillard

**Author notes:** These authors contributed equally to this work (O. Filhol-Cochet, N De Oliveira Alves, M Chamaillard). These authors share senior authorship.

## Abstract

The rapid turnover of the intestinal epithelium increases its vulnerability to genomic instability and environmental insults such as irradiation. Defects in DNA damage resolution can compromise epithelial regeneration, promote chronic tissue injury, and predispose to colorectal tumorigenesis. However, the intrinsic mechanisms that coordinate DNA repair with epithelial regeneration at the crypt level remain poorly defined. Here, we identify the Nod-like receptor protein 6 (Nlrp6) as a key epithelial regulator of genome surveillance and regenerative control in intestinal crypts. Nlrp6 is strategically expressed in crypt base columnar cells, where it preserves crypt homeostasis by restraining proliferation under genotoxic stress conditions. Loss of Nlrp6 in crypt base columnar cells results in uncontrolled oncogenic stress, defective epithelial regeneration, and accumulation of unrepaired DNA damage, features associated with poor prognosis in colorectal cancer. Conversely, aberrant Nlrp6 overexpression induces cytoplasmic retention of Csnk2 catalytic subunits, limiting their nuclear availability when DNA repair is required. These findings position Nlrp6 as a non-canonical, cell-intrinsic surveillance mechanism that links DNA damage responses to epithelial regeneration through Csnk2-dependent signaling. Collectively, our study reveals a crypt-intrinsic DNA repair pathway that governs epithelial regeneration and disease outcomes, providing new insight into how genome instability and regenerative failure contribute to colorectal cancer progression.

## INTRODUCTION

It has long been appreciated that the intestinal crypt compartment must constantly adapt to its environment for renewal of the lining of the intestine with a turnover time of 4 to 5 days in mice^1^. Every day, the proper homeostatic division of a single crypt into two daughters by a process called crypt fission is supported by small mitotically active crypt base columnar (CBC) cells. The newly differentiated intestinal epithelial cells (IECs) entirely replace existing ones through highly controlled processes involving cell multiplication, differentiation and death. The CBC are settled at the bottom of the crypt of Lieberkühn^2^, where they are strategically positioned in tiers. They are characterized by a greater expression of the leucine-rich repeat-containing G-protein-coupled receptor 5 (Lgr5), which is a member of the Wnt signaling pathway. Specifically, those bona fide active stem cells are interspersed between Paneth cells, which are discernibly specialized in secreting antimicrobial peptides and niche signals needed for regulating the balance between maintenance and differentiation^3^.

Based on lineage-tracing experiments, it has been proposed a model in which those resident stem cells double their numbers each day and stochastically adopt the fate of either a stem or transit-amplifying fate. The daughter cells that undergo functional specification will then migrate upward toward the villus tip. Perhaps more intriguing, the crypt niche is composed of additional stromal cells that remarkably instruct the fate decision of both CBC and quiescent stem cells that are located at the +4 position. To complicate matters further, the fate decision of transit-amplifying stem cells is actively adjusted by both positive and negative signals, including dietary cues and nutritional states. Notably, it has been established that fatty acid oxidation is imposed by fasting-feeding patterns for enhancing the functionality of intestinal stem cells which reside within the proliferative compartment of crypt^4^.

Along with aging, the intestinal stem cells encounter a diverse array of challenges at different time of the day which may endanger their structural integrity when they fail to repair the damage affecting their chromosomes. Given that exposure to signals derived from the gut microbiota may modulate the proliferative capacity of intestinal stem cells, we endeavored to identify which innate immune sensor may rapidly prevent from an accretion of senescent cells that may lead to oncogenic stress. The senescent phenotype consists of robust cell cycle arrest and resistance to apoptosis. It can be attained in response to numerous stressors, including DNA-damaging agents and oxidative stress. When ionizing radiation attacks host DNA, the p53 tumor suppressor gene inactivates cell cycle for DNA repair even though it remains unclear how innate immunity may prevent from neoplastic escape of senescence. It is particularly important since p53 triggers programmed cell death when DNA damage becomes irreparable^5^. The cell cycle arrest associated with senescence can deplete the pool of intestinal stem cells and transit-amplifying cells, compromising the regeneration and repair of normal intestine. unrepaired or incorrectly repaired DNA damage can lead to genomic instability and/or abnormal cell proliferation, potentially driving cancer development^6^. This led to the proposition that the attrition of stem cell numbers with oxidative stress may underlie a number of the untoward aspects of organismal aging, including the diversion to carcinogenesis.

In the present study, we found that the NOD-like receptor family pyrin domain containing 6 (Nlrp6) coordinates transcriptional program of self-renewal with crypt base columnar cells. Loss of Nlrp6 in crypt base columnar cells results in uncontrolled oncogenic stress, defective epithelial regeneration, and accumulation of unrepaired DNA damage, features associated with poor prognosis in colorectal cancer. Conversely, aberrant Nlrp6 overexpression induces cytoplasmic retention of Csnk2 catalytic subunits, limiting their nuclear availability when DNA repair is required. These findings position Nlrp6 as a non-canonical, cell-intrinsic surveillance mechanism that links DNA damage responses to epithelial regeneration through Csnk2-dependent signaling. Collectively, our study reveals a crypt-intrinsic DNA repair pathway that governs epithelial regeneration and disease outcomes, providing new insight into how genome instability and regenerative failure contribute to colorectal cancer progression.

## RESULTS

### Loss of epithelial Nlrp6 is associated with a greater persistence of DNA damage and an increased proliferative capacity of intestinal stem cells

We first examined the regeneration capacity of the intestine from Nlrp6-deficient mice after damage by 2 Gy of non-lethal ionizing radiation. At 24-hours post-irradiation, representative H&E staining of the small intestine of Nlrp6-deficient mice provided some evidence of villous blunting (Fig. 1A-B). Consistently, the number of Ki67^+^ proliferating cells per crypt was significantly decreased in Nlrp6^−/−^ mice (Fig. 1C). In parallel, the organoids from irradiated Nlrp6-deficient mice were markedly characterized by phenotypic differences when compared to controls. After 3 days in culture, proliferative cyst-like structures were more frequently observed upon loss of Nlrp6, with this distinction becoming even more pronounced by day 5 (Fig. 1D). These findings correlate with a heightened number of γ-H2AX foci within crypts (Fig. 1E and F). These data indicate a lowered capacity of intestinal stem cells to differentiate in the absence of Nlrp6. Furthermore, comparable IL-1β levels were observed in Nlrp6^−/−^ and WT mice following irradiation (Fig. 1G), indicating no significant difference in the inflammatory response. Gene ontology biological process (GOBP) analysis revealed that genes uniquely regulated in irradiated Nlrp6^−/−^ mice were enriched for humoral immune response, regulation of complement activation, and phosphatidic catabolic processes, whereas WT mice showed enrichment in rhythmic and circadian-related pathways (Fig. 1H).

**Figure 1.**
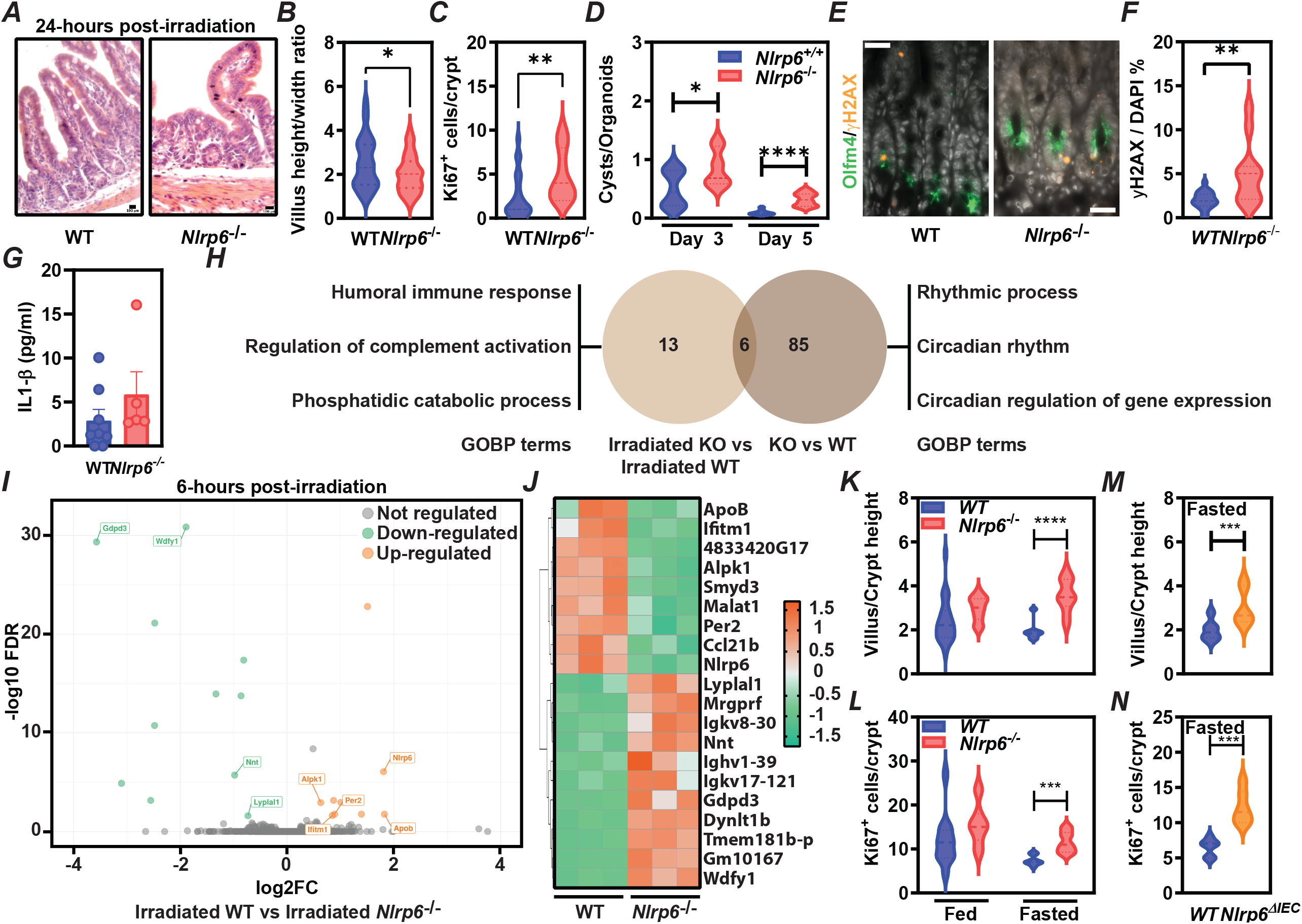
Nlrp6 deficiency exacerbates intestinal damage and alters regenerative and inflammatory responses following irradiation. (A) Representative hematoxylin and eosin (H&E)–stained sections of small intestine from WT and Nlrp6^−/−^ mice at 24 hours post-irradiation, showing altered villus architecture in the absence of Nlrp6. (B) Quantification of villus height-to-width ratio in WT and Nlrp6^−^/^−^ mice at 24 hours post-irradiation. (C) Quantification of Ki67^+^ proliferating cells per crypt, indicating reduced epithelial proliferation in Nlrp6^−^/^−^ mice. (D) Organoid formation efficiency quantified as the number of cysts/organoids derived from WT and Nlrp6^−^/^−^ intestinal crypts at days 3 and 5 post-irradiation. (E) Representative immunofluorescence images showing Olfm4 (green) and γH2AX (yellow) staining in intestinal crypts from WT and Nlrp6^−^/^−^ mice, highlighting DNA damage in stem cell compartments. (F) Quantification of γH2AX^+^ cells normalized to DAPI, demonstrating increased DNA damage in Nlrp6^−^/^−^ intestinal epithelium. (G) Measurement of IL-1β levels in intestinal tissue following irradiation, showing no significant difference between WT and Nlrp6^−/−^ mice. (H) Gene ontology biological process (GOBP) enrichment analysis of genes differentially regulated in irradiated WT versus Nlrp6^−^/^−^ mice, highlighting distinct immune- and circadian-related pathways. (I) Volcano plot depicting differentially expressed genes between irradiated WT and Nlrp6^−^/^−^ mice at 6 hours post-irradiation. (J) Heatmap showing relative expression levels of selected genes involved in metabolic, circadian, and immune processes in WT and Nlrp6^−^/^−^ mice following irradiation. (K–L) Quantification of villus/crypt height ratio (K) and Ki67^+^ cells per crypt (L) under fed and fasted conditions, indicating impaired regenerative capacity in Nlrp6^−^/^−^ mice independently of nutritional status. (M–N) Quantification of villus/crypt height (M) and Ki67^+^ cells per crypt (N) in fasted WT and Nlrp6^−^/^−^ mice at 6 hours post-irradiation. Data are presented as mean ± SEM. Statistical significance was determined using appropriate statistical tests; *p < 0.05, **p < 0.01, ***p < 0.001, ****p < 0.0001; ns, not significant. Scale bars as indicated.

In agreement with a transducing role of Nlrp6 on the ability of intestinal crypts to repopulate the damaged epithelium, bulk RNA-seq analysis revealed that loss of Nlrp6 heightened a down-regulation of several genes that are involved in mitochondrial dysfunction and in temporal maintenance of stem cells, such as Wdfy1, Nnt, Ifitm1 and Per2 (Fig. 1I and J). To conclusively establish the specific involvement of Nlrp6 on the proliferative capacity intestinal epithelial cells, we have generated a conditional knockout mice with a specific depletion of Nlrp6 in intestinal epithelial cells, including Lgr5-expresing cells. The Nlrp6Nlrp6^Δ^IEC mice carrying floxed alleles of Nlrp6 were then crossed to a strain expressing Cre recombinase under the control of the villin promoter (villin-Cre). A longer villus/crypt length and an increase in Ki67 markers were observed in the crypts of Nlrp6/Nlrp6^Δ^IEC when compared to control littermates, indicating enhanced proliferation in mice lacking Nlrp6 in intestinal epithelial cells (Fig. 1K-L). It raises the possibility that Nlrp6 signaling may enhance the differentiation for normal gut function at the expense of self-renewal capacity of intestinal stem cells. Given that short-term fasting may heighten enhance the organoid-forming capacity of intestinal stem cells, we asked whether the shift from dormant state to a state of self-renewal may depend on NLRP6 expression after exposure to ionizing radiation. While the villus/crypt ratio exhibited a significant increase as early as 18 hours earlier compared to the group expressing this protein and enhanced proliferation in the crypts of Nlrp6-deficient mice, particularly under fasting conditions (Fig. 1M and N).

### Strategic function of Nlrp6 in Lgr5^+^ intestinal stem cells prevent colorectal cancer initiation

Having determined that the presence or absence of Nlrp6 within intestinal epithelial cells may modulates the capacity of intestinal stem cells to repopulate the damaged epithelium, we next decided to sort intestinal stem cells by fluorescence-activated cell sorting (FACS) from the colon of Lgr5-EGFP-ires-creERT2 mice (Supplementary Fig.1), which express the enhanced green fluorescent protein (eGFP) under the control of the Lgr5 promoter. Using publicly available single-cell RNA-seq data, single-cell transcriptomic analysis revealed that Nlrp6 expression is highly enriched in Lgr5^+^ intestinal stem cells, with comparatively lower expression in differentiated epithelial lineages and immune cell populations (Fig.2A). In addition, transcriptional RNA profiling of sorted Lgr5+ cells by an unbiased bulk population RNA sequencing (RNA-seq) analysis allowed us to systematically examine the expression of all innate immune sensors. It revealed that intestinal stem cells may not express a broad (but rather a limited) portfolio of Toll-like receptors and Nod-like receptors. Among those, the transcript level of the gene encoding for Nlrp6 was by far the most abundant in continuously cycling intestinal stem cells at steady-state (Fig. 2B).

**Figure 2.**
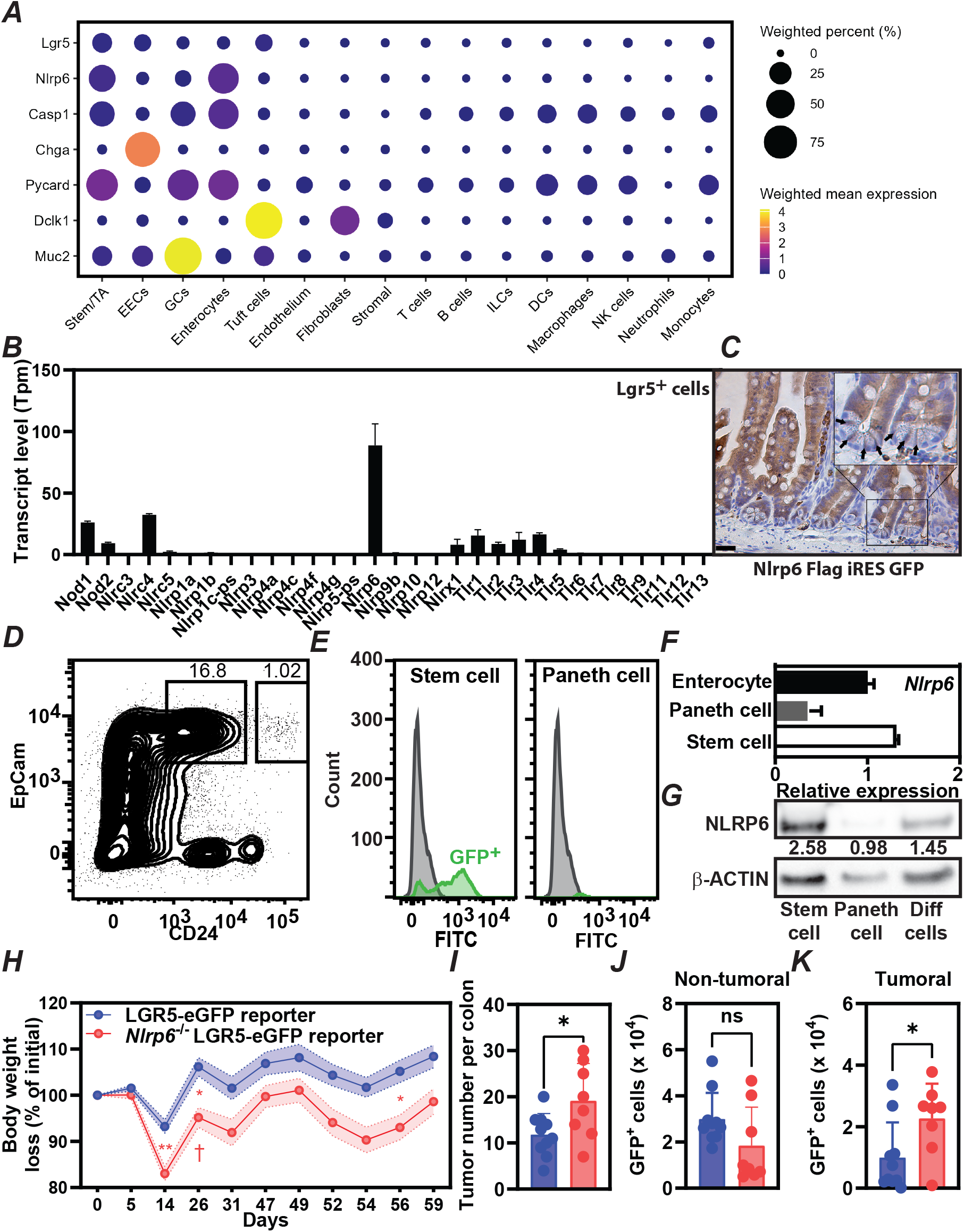
Nlrp6 is selectively expressed in Lgr5^+^ intestinal stem cells and restrains colorectal tumor initiation. (A) Dot plot showing single-cell RNA-sequencing expression of Lgr5, Nlrp6, inflammasome-related genes, and epithelial markers across intestinal cell populations. Dot size indicates the percentage of expressing cells and color intensity reflects mean expression levels. (B) Transcript levels (TPM) of NLR and TLR family members in sorted Lgr5^+^ intestinal stem cells. (C) Immunohistochemical staining of intestinal tissue from Nlrp6-Flag-IRES-GFP reporter mice showing Nlrp6 expression localized to crypt base stem cells. (D–E) Flow cytometric identification of EpCam^+^CD24^low stem cells and GFP expression in stem, Paneth, and enterocyte populations. (F) Relative Nlrp6 mRNA expression in sorted intestinal epithelial populations. (G) Immunoblot analysis of NLRP6 protein levels in stem cells, Paneth cells, and differentiated epithelial cells, with β-actin as loading control. (H) Body weight loss over time in WT and Nlrp6^−^/^−^ Lgr5-eGFP reporter mice during tumorigenesis. (I) Quantification of tumor number per colon. (J–K) Quantification of GFP^+^ stem cells in non-tumoral (J) and tumoral (K) tissues. Data are presented as mean ± SEM; *p < 0.05; **p < 0.01; ns, not significant.

To validate this, we generated a knock-in mouse expressing Nlrp6 with a carboxy-terminal 3xFlag Tag and an IRES-eGFP cassette (Supplementary Fig. 2A and B). We then examined the pattern of Nlrp6 expression at the protein level using immunohistochemistry and immunofluorescence analysis of both small and large intestine of *Nlrp6-FLAG-ires-EGFP* knock-in mice (Fig. 2C). FACS analysis confirmed that Nlrp6 is reliably expressed by intestinal stem cells and their transit-amplifying daughters when compared to that in post-mitotic Paneth cells where transcript levels of Nlrp6 were substantially lower (Fig. 2D and E). Accordingly, the antibody against either GFP or Nlrp6 detected a protein product migrating at ∼100kDa, which is consistent with the predicted size of Nlrp6 (Fig. 1F). Densitometric analysis of western-blots confirmed that Nlrp6 protein is expressed in intestinal stem cells to the same extent as what is observed in enterocytes. Even though this does not exclude the possibility that NLRP6 may regulate Paneth cells function in nurturing stem cells, we failed to observe protein level of Nlrp6 in Paneth cells if any (Fig. 2G). This restricted pattern of Nlrp6 expression to Lgr5^+^ intestinal stem cells suggest its contribution to their renewal.

To assess the functional relevance of Nlrp6 in Lgr5^+^ stem cells during colorectal tumorigenesis, Nlrp6^−/−^ Lgr5-eGFP reporter mice were subjected to tumor induction. Nlrp6-deficient mice exhibited increased body weight loss during disease progression compared with WT controls (Fig. 2H). Importantly, loss of Nlrp6 resulted in a significantly higher tumor burden per colon (Fig. 2I). While the number of GFP^+^ stem cells in non-tumoral tissue was not significantly altered (Fig. 2J), tumoral regions from Nlrp6^−^/^−^ mice displayed a marked expansion of GFP^+^ cells (Fig. 2K), indicating enhanced stem cell accumulation within tumors. Collectively, these results demonstrate that Nlrp6 is selectively expressed in Lgr5^+^ intestinal stem cells and functions as a critical regulator limiting stem cell–driven colorectal cancer initiation and progression.

### The proliferative capacity of transit-amplifying cells is heightened in the absence of Nlrp6 at steady-state

Since genetic ablation of Nlrp6 is causing a greater expression of Wnt-target genes^7^, we next decided to define to which extent the properties of Lgr5-expressing intestinal stem cells is altered upon loss of Nlrp6. To this end, Lgr5^+^ cells were sorted from the colon of Lgr5-EGFP mice that were crossed or not with Nlrp6^−/−^ mice. The gene expression profiling of Lgr5^+^ intestinal stem cells was then examined through processing of bulk-population RNA sequencing data under steady state conditions. Supporting a role for Nlrp6 on the proliferation capacity of stem cell function, computational analysis revealed an over-representation of differentially expressed genes that are linked to the regulation of TLR signaling and negative regulation of Wnt signaling. Key Wnt-associated genes such as Cav1, Wwtr1, and Shl3 are dysregulated in Nlrp6^−/−^ cells, indicating that Nlrp6 modulates Wnt pathway homeostasis in the intestinal epithelium (Fig. 3A and B).

**Figure 3.**
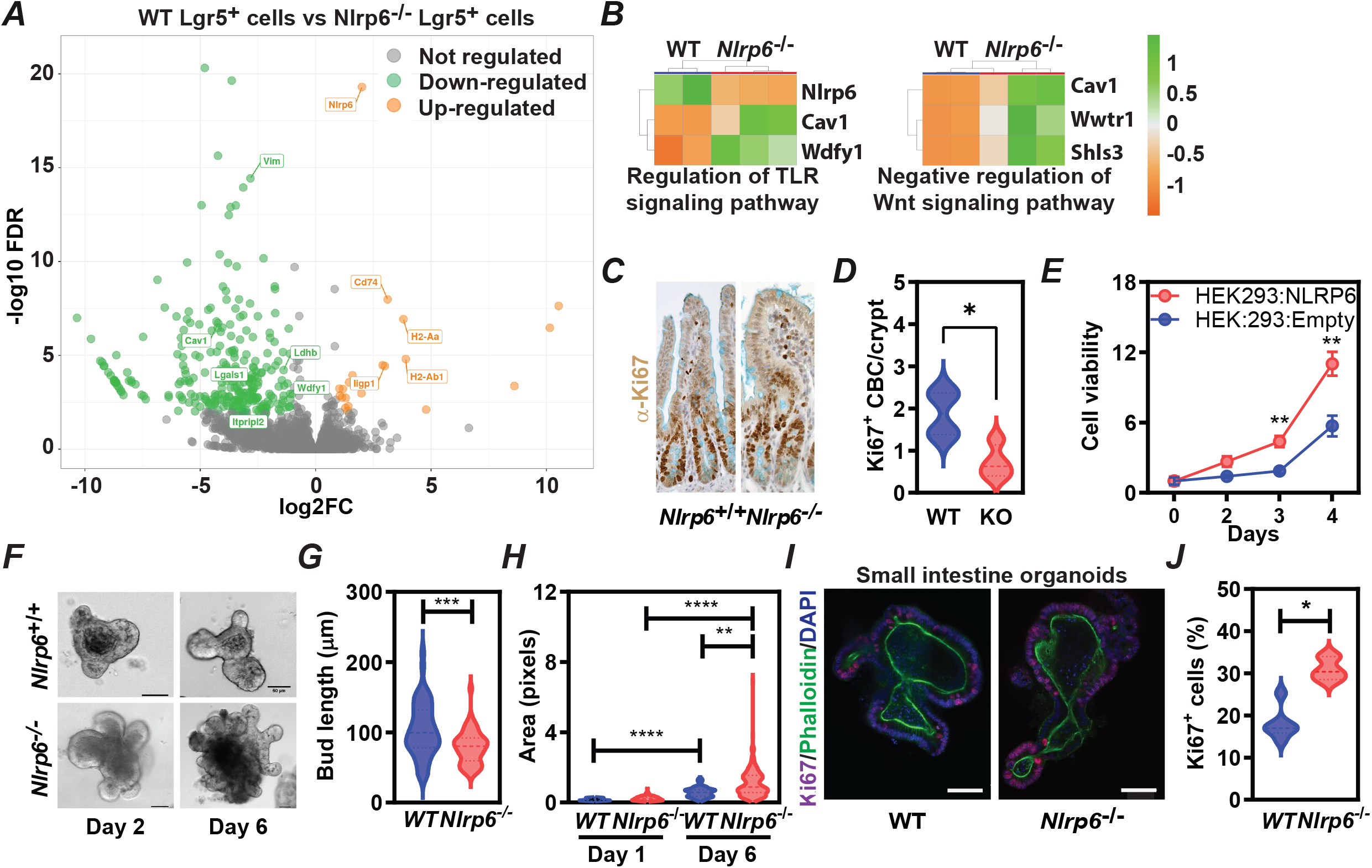
Loss of Nlrp6 in Lgr5^+^ intestinal stem cells enhance epithelial proliferation. (A) Differential transcriptomic analysis comparing WT and Nlrp6^−/−^ Lgr5^+^ intestinal stem cells. The volcano plot shows a broad transcriptional reprogramming in the absence of Nlrp6, with a substantial number of upregulated genes. Several differentially expressed genes are associated with epithelial remodeling and Wnt-related pathways, suggesting that Nlrp6 restrains pro-proliferative transcriptional programs in intestinal stem cells. (B) Functional enrichment and pathway analysis of differentially expressed genes highlight significant alterations in innate immune signaling and negative regulation of Wnt signaling. (C–D) Immunohistochemical staining for Ki67 within intestinal crypts of Nlrp6^−^/^−^ mice compared with WT controls. (E) Expression of Nlrp6 in HEK293 cells significantly reduces cell viability over time, supporting a cell-intrinsic role for NLRP6 in limiting cellular growth and proliferative capacity. (F–H) intestinal organoids derived from Nlrp6^−^/^−^ mice display pronounced morphological alterations, including increased size and area compared with WT organoids. (I–J) Immunofluorescence analysis of small intestinal organoids shows a significant increase in the proportion of Ki67^+^ cells in Nlrp6^−^/^−^ organoids relative to WT.

Given that Wnt signaling is the central driver of intestinal stem cell maintenance and proliferation^8^, we examined cell cycle progression through Ki67 antibody staining, a marker expressed in all cell cycle phases. Intriguingly, the loss of Nlrp6 expression resulted in an enhanced proportion of Ki67^+^ cells in transit-amplifying cells at the expense of the long-lived pool of cycling stem cells (Fig. 3C). Quantification confirms a significant expansion of Ki67^+^ crypt base columnar cells, demonstrating that loss of Nlrp6 enhances epithelial proliferation *in vivo* (Fig. 3D). Consistently, expression of NLRP6 in HEK293 cells significantly reduces cell viability over time, supporting a cell-intrinsic role for NLRP6 in limiting cellular growth and proliferative capacity (Fig. 3E).

Microscopy analysis of three-dimensionally cultured cyst-like structures showed a greater expression of Nlrp6 at the tips of organoid buds in control (Fig. 3F and G). This was further corroborated by a greater number of Ki67^+^ cells led to a greater average area of the Nlrp6-deficient organoids than controls, with this distinction becoming even more pronounced by day 6 (Fig. 3H), indicating that intestinal stem cells could survive in the absence of Nlrp6. In line with these findings, immunofluorescence analysis of small intestinal organoids shows a significant increase in the proportion of Ki67^+^ cells in Nlrp6^−/−^ organoids relative to WT (Fig. 3I and I). Together, these data establish Nlrp6 as a critical negative regulator of intestinal epithelial proliferation, acting to constrain stem and progenitor cell expansion both *in vivo* and *ex vivo*.

### Nlrp6 interacts with the regulatory subunit of Csnk2 to limit intestinal epithelial proliferation

We then set out to understand the specific molecular machinery by which Nlrp6 may drive this phenomenon. We performed a bacterial two-hybrid screening of a MEFs cDNA library by applying the Pyrin domain of mouse Nlrp6 as bait. Two positive overlapping clones contained a cDNA encoding for the regulatory beta subunit of Phosvitin/casein kinase type II (encoded by Csnk2β, OMIM 115441). Csnk2 is a highly conserved tetrameric serine/threonine-selective protein kinase that is evolutionarily conserved throughout the eukaryotic kingdom. It forms a heterotetramer composed of two catalytic subunits (Csnka1 and Csnka2, α and α’ subunits) and a dimer of regulatory subunits (Csnkb1 and Csnkb2, β subunits), which are found in both the nucleus and cytoplasm. It has emerged that this protein kinase plays an essential role in the DNA repair process by monitoring and repairing both single and double-strand breaks^9^. As predicted by our two-hybrid screening, Nlrp6 endogenously immunoprecipitated with Csnk2β in the mouse intestine (Fig. 4A). Endogenous Nlrp6 co-precipitated with Csnk2β in wild-type samples but not in Nlrp6^−/−^, confirming the specificity of the interaction (Fig. 4B).

**Figure 4.**
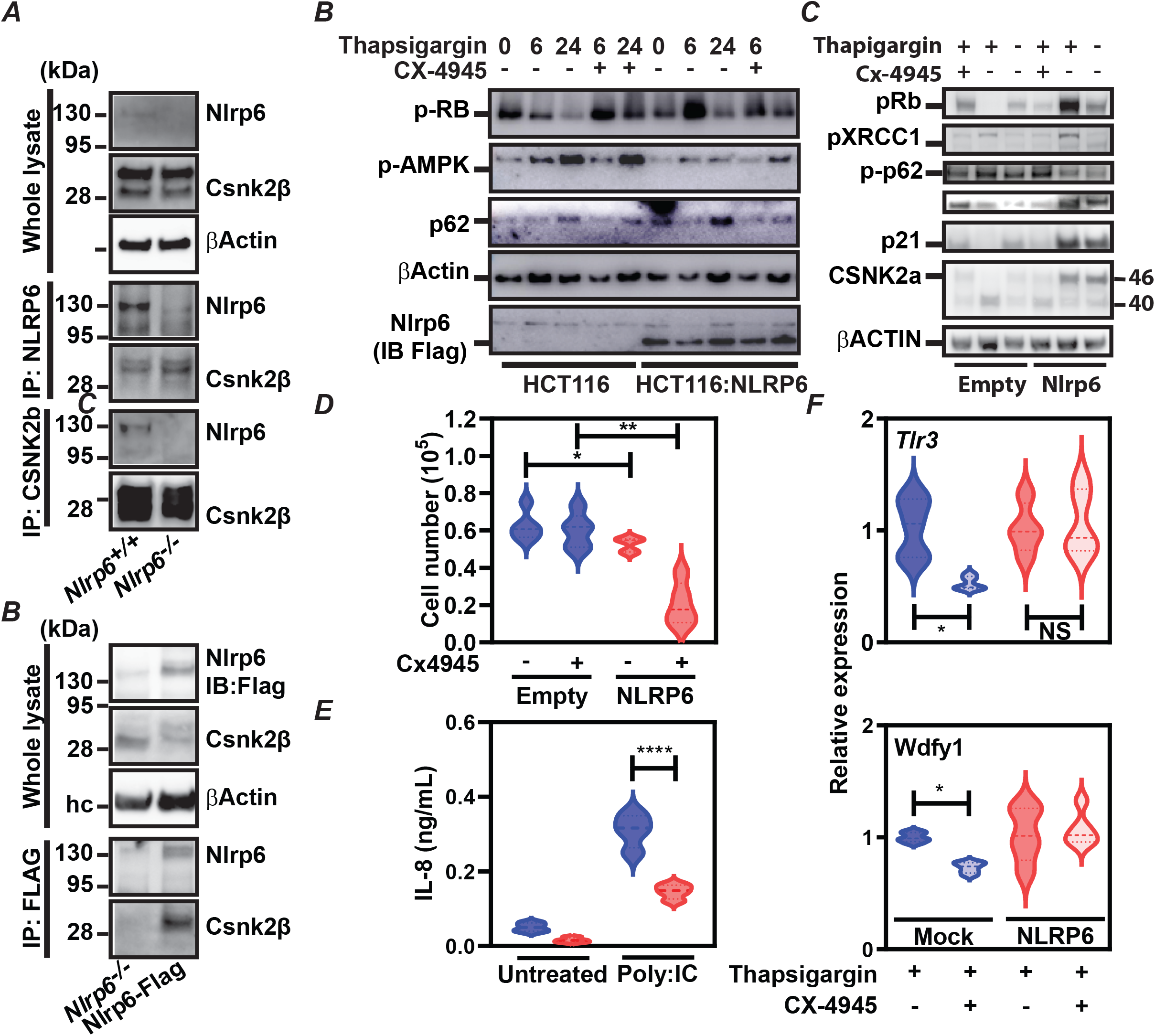
NLRP6 interacts with CSNK2β and limits proliferative and stress-response signaling. (A) Co-immunoprecipitation analysis showing the interaction between NLRP6 and CSNK2β in Nlrp6^+/+^ and Nlrp6^−/−^ cells. Whole-cell lysates and immunoprecipitates (IP: NLRP6 or IP: CSNK2β) were analyzed by immunoblotting as indicated. (B) Immunoblot analysis of HCT116 and HCT116-NLRP6 cells treated with thapsigargin in the presence or absence of CX-4945 for the indicated time points. Levels of phosphorylated Rb (p-RB), p-AMPK, p62, and NLRP6 (IB: Flag) were evaluated. β-Actin was used as a loading control. (C) Immunoblot analysis of cells expressing empty vector or NLRP6 following treatment with thapsigargin and/or CX-4945. Expression of pRb, pXRCC1, p-p62, p21, and CSNK2α was assessed. (D) Quantification of cell numbers under the indicated conditions in empty vector and NLRP6-expressing cells treated with or without CX-4945. Data are presented as mean ± SD. p values are indicated. (E) IL-8 secretion (ng/mL) measured in untreated or Poly(I:C)-treated cells expressing NLRP6. (F) Relative expression of Tlr3 and Wdfy1 under mock or NLRP6-expressing conditions with thapsigargin and/or CX-4945 treatment.

To assess the functional consequences of this interaction, HCT116 control and NLRP6-expressing HCT116 cells were treated with the endoplasmic reticulum stress inducer thapsigargin in the presence or absence of the Csnk2 inhibitor CX-4945. Immunoblot analysis showed that Nlrp6 expression attenuated thapsigargin-induced phosphorylation of Rb and Ampk, while modulating p62 accumulation, effects that were reversed by Csnk2 inhibition at 24h (Fig. 4C and D). Consistently, Nlrp6 expression reduced cell proliferation, as evidenced by a significant decrease in cell numbers following CX-4945 treatment compared with empty vector controls (Fig. 4E).

Functionally, Nlrp6 expression significantly reduced inflammatory responsiveness, as poly(I:C)-induced IL-8 secretion was markedly reduced in Nlrp6-expressing cells relative to controls (Fig. 4F). Finally, transcriptional analysis demonstrated that Nlrp6 modulated stress-associated gene expression, with significant regulation of Tlr3 and Wdfy1 expression following thapsigargin and CX-4945 treatment, indicating that Nlrp6-Csnk2 signaling influences epithelial stress adaptation pathways (Fig. 4G). Together, these data support a model in which Nlrp6 constrains epithelial proliferation and stress signaling by modulating Csnk2-dependent pathways.

## DISCUSSION

Maintenance of intestinal health and regeneration require precise coordination among proliferation, differentiation, and genome surveillance to preserve tissue integrity while enabling rapid repair^10^. Here, we identify Nlrp6 as a central cell-intrinsic regulator of this balance, acting specifically within Lgr5^+^ intestinal stem cells to restrain excessive proliferation, promote differentiation, and safeguard genomic stability under both homeostatic and stress conditions.

A striking observation of this study is that loss of epithelial Nlrp6 results in persistent DNA damage following ionizing radiation, as evidenced by sustained γ-H2AX foci, despite comparable levels of IL-1β. This dissociation between DNA damage accumulation and inflammation challenges the prevailing view of Nlrp6 as a direct regulator of epithelial stress responses. Persistent DNA damage in intestinal stem cells is known to favor clonal expansion, aberrant repair, and tumor initiation^11^, suggesting that Nlrp6 functions as a molecular safeguard limiting the propagation of genomically unstable epithelial cells.

Our data further reveal that Nlrp6 regulates the balance between proliferation and differentiation in the intestinal epithelium. While Nlrp6 deficiency enhances crypt cell proliferation and organoid growth, it concomitantly impairs differentiation, leading to cyst-like organoid structures and altered crypt-villus architecture. The loss of coordination between proliferation and differentiation represents a key early event in tumor development, reflecting altered stem cell homeostasis^12^. We propose that Nlrp6 constrains self-renewal programs to preserve epithelial organization, even at the expense of maximal regenerative output.

Transcriptomic analyses further uncovered an unexpected link between Nlrp6 and circadian and metabolic regulation in intestinal stem cells. Interestingly, irradiated colon samples from Nlrp6-deficient mice exhibited dysregulation of genes involved in mitochondrial redox balance (Nnt), alongside alterations in pathways controlling membrane dynamics and lipid remodeling, including Wdfy1, Lypla1, and Gdpd3. Wdfy1, a FYVE-domain–containing adaptor implicated in endosomal trafficking and TLR3/4 signaling, has been linked to innate immune activation and vesicular sorting processes^13^. Lypla1, a lysophospholipase and depalmitoylating enzyme, regulates dynamic protein S-palmitoylation cycles that control membrane association and signal transduction^14^. Notably, suppression of LYPLA1 has been shown to inhibit cell proliferation and migration in non-small cell lung cancer cells, underscoring its role in oncogenic signaling and membrane-associated growth pathways^15^.

The relevance of Nlrp6 in adaptive stem cell plasticity is further highlighted by fasting experiments. Short-term fasting, a condition known to transiently enhance ISC function through metabolic rewiring^16^, markedly amplified crypt proliferation in Nlrp6-deficient mice following irradiation. These findings suggest that Nlrp6 preventing excessive or prolonged proliferative responses under metabolic stress. In Nlrp6-deficient conditions, adaptive programs may become dysregulated, compromising fundamental cellular control mechanisms.

At the cellular level, we demonstrate that Nlrp6 expression is selectively enriched in Lgr5^+^ ISCs, with minimal expression in differentiated epithelial lineages. This restricted expression pattern strongly implicates Nlrp6 as a stem cell–specific regulator rather than a general epithelial sensor. Previous studies have linked Nlrp6 to intestinal homeostasis, epithelial integrity, with emerging evidence indicating that its loss can exacerbate inflammation-driven tumorigenesis ^17,18^. Here, our data extend this framework by positioning Nlrp6 directly within the stem cell compartment as a stem cell-specific tumor-suppressive checkpoint. Thus, rather than acting as a general epithelial sensor, Nlrp6 appears to function as a selective barrier that limits malignant progression at the level of stem cell hierarchy.

Mechanistically, our identification of Csnk2 as an endogenous Nlrp6 interactor uncovers a surveillance pathway that restrains oxidative stress-associated damage and safeguards against colorectal cancer proliferation. Csnk2 has emerged as a key participant in the DNA repair process, essential for monitoring and repairing both single and double-strand breaks ^9,19^. For instance, it was proposed that Csnk2 inhibition triggers cell death by early disruption of mitochondrial Ca^2+^ dynamics, altering mitochondrial permeability and membrane potential^20^. Here, Nlrp6 modulates Csnk2-dependent phosphorylation of key regulators such as Rb and AMPK and attenuates stress-induced proliferative signaling. These effects are reversed by pharmacological inhibition of Csnk2, indicating that Nlrp6 constrains epithelial proliferation through Csnk2-dependent stress-adaptation pathways. This positions Nlrp6 as a molecular brake that limits cell cycle progression when genomic or metabolic stress remains unresolved.

Collectively, our findings redefine Nlrp6 as a multifunctional regulator of intestinal stem cell fate, integrating genome surveillance, metabolic adaptation and kinase signaling to maintain epithelial homeostasis. Loss of this regulatory axis uncouples regeneration from differentiation and DNA repair, creating a permissive environment for stem cell–driven colorectal cancer initiation. Taken together, these data suggest that innate immune sensors may have evolved roles beyond host defense, acting as intrinsic guardians of stem cell integrity in rapidly renewing tissues.

## MATERIALS AND METHODS

### Cell lines

Mouse embryos were obtained from two Nlrp6^+/+^ and two Nlrp6^-/-^ pregnant mice after close to 15 days and 13.5 days of gestation for Nlrp6^+/+^ and Nlrp6^-/-^, respectively, and mouse embryonic fibroblasts (MEF) were cultured in accordance with the 3T3 protocol. Specifically, two embryonic fibroblasts cultures were produced in parallel, each from a pool of cells obtained from four embryos, from each WT mouse, for a total of four 3T3 MEF cells cultures in parallel. Similarly, two embryonic fibroblasts cultures were produced in parallel, each from a pool of cells obtained from seven embryos, from each Nlrp6^-/-^ mouse, for a total of four 3T3 MEF cells cultures in parallel. Briefly, cells were passaged every three days by seeding a fixed amount of cells in a new cell culture flask, and cell count by trypan blue exclusion was recorded at each passage, allowing the detection of growth speed decrease and entry into senescence by calculation of the Passage Doubling Level (PDL). To calculate cumulative population doublings, the following formula was applied: PDL = 3.32 (log Xe − log Xb) + S, where Xb = cell number at the beginning of the incubation time, Xe = the cell number at the end of the incubation time, and S is the starting PDL of the inoculum. The mean Cumulative PDL during the first days of culture until reaching senescence was plotted with standard error of the mean (s.e.m.) for n=4 cultures in parallel for Nlrp6^+/+^ and Nlrp6^-/-^ genotypes.

### Single-cell gene expression analysis of inflammasome components

Single-cell RNA sequencing (scRNA-seq) data from colon samples of adult wild-type C57BL/6Ntac mice were retrieved from the Broad Institute Single Cell Portal under accession number SCP2760. The dataset was subset to analyze a specific panel of inflammasome-related genes: Lgr5, Nlrp6, Casp1, Chga, Pycard, Dclk1, and Muc2. Cells were aggregated by broad lineage identity based on the original annotations. For each lineage, the percentage of expressing cells and the mean expression levels were calculated. Visualization was performed using ggplot2 in R software.

### RNAseq analysis and Gene set enrichment analysis

Lgr5-expressing stem cells and mice colon tissue were lysed in RLT buffer and RNA was extracted using RNeasy Mini Kit (QIAGEN) following manufacturer protocol. RNA quality was checked using a Bioanalyzer 2100 (Agilent technologies). mRNA library preparation was realized following the manufacturer’s recommendations (Illumina Stranded mRNA Prep Kit from ILLUMINA). Final samples pooled library prep was sequenced on Novaseq 6000 ILLUMINA with S1-200cycles cartridge (2x1600Millions of 100 bases reads) corresponding to 2x30Millions of reads per sample after demultiplexing. Differentially expressed genes (DEGs) were identified with DESeq2 method and further analyzed using over-representation analysis from the Molecular Signatures Database (MSigDB) database.

### Immunoblotting analysis

Cells or tissue samples were harvested in RIPA Lysis buffer with protease inhibitor (Roche, Basel, Switzerland) and phosphatase inhibitor (Sigma-Aldrich, St.Louis, MO). Total protein extracts were quantified using the Pierce BCA Protein Assay kit (Thermo Scientific, USA). Five micrograms of total protein from cell extracts were separated on a 4-12% SDS polyacrylamide gel, and blotted onto a nitrocellulose membrane. Membranes were blocked for 1 h in 5% BSA in PBS-Tween 20 (0.05%) (PBST), incubated overnight at 4 °C with primary antibody, washed three times with PBST, and incubated for 1 h with secondary antibody. After final washes with PBST, the blots were developed using an enhanced chemiluminescence detection system (Pierce™ ECL Substrate or SuperSignal™ West Femto, Thermo Fisher) and analysis was performed using the primary antibody and appropriate HRP-conjugated secondary antibody.

### Immunohistochemistry and immunofluorescence staining

Ileum samples were fixed in 4% neutral formaldehyde, dehydrated, paraffin-embedded, sliced into 5 μm thick sections using a Leica RM 2155 microtome. These sections were then used for histological slide preparation, stained with hematoxylin and eosin, and analyzed under an optical microscope. Rehydration of the samples involved sequential immersion in specific solutions: Toluene (twice for 3 minutes each), 100% Ethanol (twice for 3 and 1 minute respectively), 95% Ethanol (1 minute), 70% Ethanol (1 minute), and Milli-Q water (1 minute). Antigen retrieval was performed by incubating the deparaffinized tissues in a 10 mM sodium citrate solution (pH 6) for 20 minutes at 95°C, followed by a 5-minute PBS bath. To enhance specificity, endogenous peroxidases were inactivated by adding 25 µL of 3% hydrogen peroxide for 10 minutes in a dark, humidified environment at room temperature, after delineating tissue squares using the ImmEdge™ pen. Sections were incubated in blocking buffer (PBS, 5% BSA, 0.3% Triton X-100) for 1h at RT.

For ki67 immunostaining, the sections were incubated overnight in a humidified, dark chamber with 25 μL of anti-Ki67 antibody (diluted 1:750). Sections were washed three times using PBS for 5 min at RT and the appropriate HRP-conjugated secondary antibody was incubated in antibody diluent for 1h at RT. Finally, visualization was achieved by adding 25 μL of DAB-substrate for 5 minutes. Villus and crypt length (μm) of ileal tissue were measured and the number of Ki67-positive cells was counted from crypts per mouse per group.

For immunofluorescence, primary antibodies were incubated overnight at 4°C in antibody diluent. Sections were washed three times using PBS for 5 min at RT and incubated with corresponding secondary antibodies diluted in antibody diluent for 1h at RT. Slides were washed thrice in PBS and nuclei were stained with DAPI solution (1 μg/ml) for 3 min at RT and mounted with ProLong Gold antifade mounting solution (ThermoFisher Scientific). The following primary antibodies were used: anti-H2A.X Phospho (Ser139); anti-Olfm4, p-XRCC1.

### RNA extraction and quantitative real-time PCR (qRT-PCR) from mice samples

Ileum tissues were homogenized with ceramic beads on a MagNA Lyser (Roche). RNA was extracted using RNeasy Mini Kit (QIAGEN) following manufacturer protocol. 250ng RNA of each sample was retro-transcribed using Affinity Script cDNA synthesis kit (Agilent Technologies). The resulting cDNA (equivalent to 5 ng of total RNA) was amplified using the SYBR Green real-time PCR kit and detected on either MxPro or AriaMx (Agilent Technologies). qRT-PCR analysis was performed with the forward and reverse primers that were designed using Primer 3 software (sequences available upon request). On completion of the qPCR amplification, a DNA melting curve analysis was carried out to confirm the presence of a single and specific amplicon. Actb were used as an internal reference gene to normalize the transcript levels of each gene of interest. Relative mRNA levels (2-2-ΔΔCt) were determined by comparing (i) the PCR cycle thresholds (Ct) for Actb and the genes of interest (ΔCt); (ii) ΔCt values for treated and control groups (ΔΔCt).

### Cell proliferation assay

Proliferating cells were detected at various times with the WST1 cell-proliferation assay (ab155902) and were measured according to the manufacturer’s instructions. The absorbance was measured using FLUOstar Omega plate reader (BMG Labtech) at 420 nm for each time point.

