## Supplementary figures and images for "A crypt-operating DNA repair checkpoint for uncoupling regeneration and tumorigenesis"

### Supplementary Fig.1

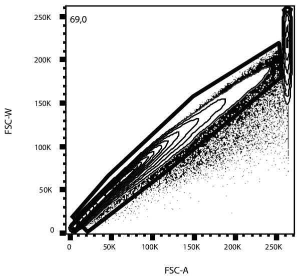


7075.fcs
Ungated
160271

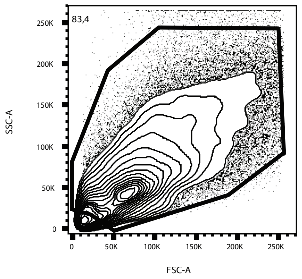


7075.fcs
Single
110616

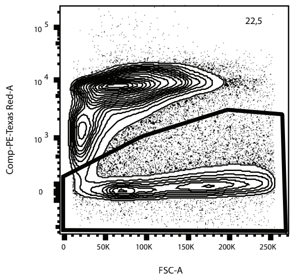


7075.fcs
Cells
92248

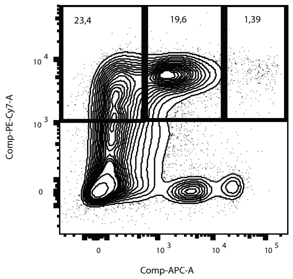


7075.fcs
Live
20778

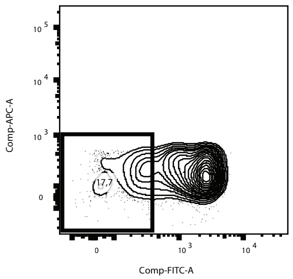


7075.fcs
Epcam+ CD24 low
4857

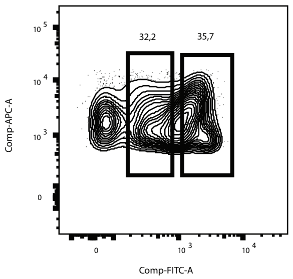


7075.fcs
Epcam+ CD24 med
4064

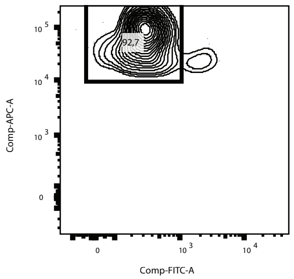


7075.fcs
Epcam+ CD24 high
288

### Supplementary Fig.2

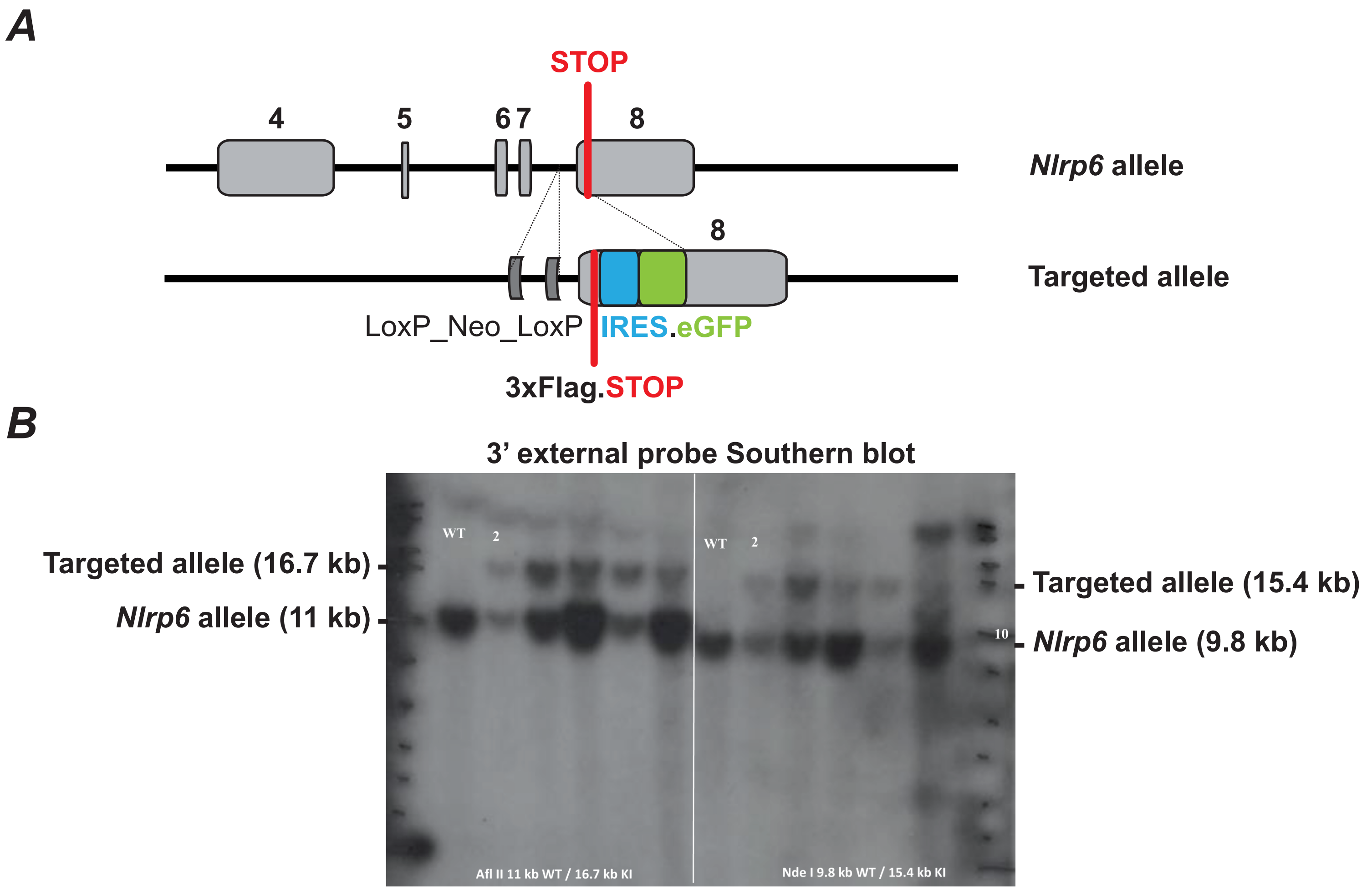
